# Spindle Assembly Checkpoint Competency Determines Sensitivity to KIF18A Inhibition in Small-Cell Lung Cancer

**DOI:** 10.64898/2026.07.30.741836

**Authors:** Chiori Tabe, Rajesh Kumar, Yue Huang, Michael J. Kruhlak, Ajit K. Sharma, Roshan L. Shrestha, Anish Thomas

## Abstract

**Background:** Small-cell lung cancer (SCLC) is characterized by pervasive chromosomal instability (CIN) and remains largely refractory to targeted therapies. KIF18A, a motor protein that regulates chromosome alignment during mitosis, has emerged as a selective dependency in CIN-high tumors. Whether this dependency extends to SCLC, a prototypical CIN-high cancer, has not been established, and biomarkers predicting response to KIF18A inhibition, currently in clinical trials, are lacking.

**Methods:** We integrated analyses of patient tumor datasets, neuroendocrine (NE) and non- NE SCLC cell lines, and functional perturbation models to define the determinants of response to KIF18A inhibition. Chromosomal instability metrics, transcriptional programs, mitotic dynamics, and spindle assembly checkpoint (SAC) function were assessed using genomic profiling, live-cell imaging, genetic perturbation, and pharmacologic inhibition.

**Results:** KIF18A expression was elevated in SCLC tumors and correlated with CIN-associated transcriptional programs, proliferative markers, and NE status; however, these features did not predict sensitivity to KIF18A inhibition. Instead, response was determined by the functional integrity of the SAC. SAC-proficient SCLC cells underwent sustained mitotic arrest followed by apoptotic cell death upon KIF18A inhibition, whereas SAC-defective cells failed to maintain checkpoint activation and survived. Mechanistically, resistant cells exhibited impaired kinetochore recruitment of core SAC components, including MAD1 and BUBR1. Importantly, transient induction of acute CIN through MPS1 inhibition partially restored sensitivity to KIF18A inhibition in resistant models.

**Conclusions:** This study provides the first mechanistic characterization of KIF18A dependency in SCLC, identifying SAC competency as the primary determinant of response. These findings establish a biologically informed framework for patient stratification and rational combination strategies.

**Translational Relevance:** Small-cell lung cancer (SCLC) is an aggressive malignancy with few effective targeted therapies and marked chromosomal instability. KIF18A has emerged as a potential therapeutic target in genomically unstable cancers, but biomarkers predicting response to KIF18A inhibition are lacking. We demonstrate that sensitivity to KIF18A inhibition in SCLC is determined not by KIF18A expression, neuroendocrine subtype, or baseline chromosomal instability, but by the functional integrity of the spindle assembly checkpoint (SAC). SCLC cells with intact SAC signaling undergo sustained mitotic arrest and apoptosis upon KIF18A inhibition, whereas SAC-defective cells bypass checkpoint activation and survive aberrant mitosis. Notably, transient induction of acute chromosomal instability through MPS1 inhibition partially restores sensitivity in resistant models. Together, these findings identify mitotic checkpoint competency as a mechanistic determinant and candidate predictive biomarker for KIF18A-targeted therapies, providing a biologically informed framework for patient stratification and rational combination strategies relevant to ongoing KIF18A inhibitor clinical trials.

## Introduction

Small-cell lung cancer (SCLC) is a highly aggressive neuroendocrine carcinoma characterized by rapid disease progression, early metastatic dissemination, and poor prognosis (1). Although initially sensitive to chemotherapy, most patients relapse within months, and the 5-year survival rate remains below 7% (2). Genomic profiling has revealed near-universal inactivation of TP53 and RB1 and widespread genomic rearrangements, distinguishing SCLC from non-small-cell lung cancer and highlighting the paucity of recurrent, targetable oncogenic drivers (3). As a result, therapeutic advances have been limited for decades, underscoring the urgent need to identify novel vulnerabilities that can be exploited for durable clinical benefit.

Chromosomal instability (CIN) is a pervasive feature of many cancers and has been implicated in driving malignant progression, intratumoral heterogeneity, and therapy resistance (4,5). SCLC is defined by exceptionally high proliferative rates, which drive profound CIN (6). Indeed, replication stress arising from rapid cell-cycle progression has been mechanistically linked to both numerical and structural CIN, contributing to the mitotic aberrations characteristic of fast-cycling tumors (7,8). Importantly, chromosomally unstable cells often rely on specific mitotic regulators to maintain genome integrity and viability, revealing potential synthetic-lethal vulnerabilities (9). Despite the extensive genomic instability that defines SCLC (3,10), the therapeutic potential of targeting mitotic dependencies in this disease remains largely unexplored.

SCLC exhibits marked transcriptional and phenotypic heterogeneity. Distinct molecular subtypes driven by ASCL1, NEUROD1, POU2F3, and YAP1 differ in neuroendocrine differentiation, proliferative behavior, metabolic programs, and therapeutic response (11). Moreover, intratumoral heterogeneity and lineage plasticity enable dynamic transitions between states, particularly under treatment pressure, complicating patient stratification and biomarker development (12). As a result, vulnerabilities inferred from static genomic features may not uniformly translate across SCLC subtypes or cellular states, underscoring the need for functionally defined dependencies.

Faithful chromosome segregation during mitosis depends on the coordinated activity of microtubule and kinetochore-associated, and mitotic checkpoint complex-associated proteins that function to monitor and maintain the integrity of the spindle assembly checkpoint (SAC). KIF18A, a member of the kinesin-8 family is a molecular motor that regulates microtubule dynamics to promote accurate chromosome alignment and segregation (9,13,14). Loss or inhibition of KIF18A leads to persistent chromosome misalignment, activation of the SAC, prolonged mitotic arrest, and subsequent cell death (13,15). Prior studies have demonstrated that diploid or genomically stable cells tolerate loss of KIF18A functions, whereas it is selectively essential for cancer cells with high CIN, where its inhibition results in mitotic delay and lethality (9,15–17). These findings suggest that KIF18A serves as a compensatory mechanism that enables CIN-high cells to survive under conditions of persistent mitotic stress.

Pharmacologic targeting of KIF18A has recently become feasible. ATP-competitive inhibitors such as AM-9022, along with other emerging KIF18A-directed compounds, have demonstrated preclinical efficacy in CIN-high tumor models by disrupting spindle dynamics and inducing SAC-dependent mitotic arrest (17). Importantly, multiple KIF18A inhibitors have now advanced into early-phase clinical trials in patients with solid tumors, underscoring the translational relevance of mitotic dependency. Given the paucity of targetable oncogenic drivers and the high degree of CIN, targeting KIF18A could offer a novel therapeutic approach to selectively eliminate vulnerable SCLC subpopulations. Defining the molecular determinants of sensitivity and resistance will be essential to guide patient selection and rational combination strategies as these agents move forward clinically.

Here, we investigate the role of KIF18A in SCLC, assess the sensitivity of molecularly distinct SCLC models to a selective KIF18A inhibitor, and delineate the mechanisms underlying differential drug responses. Our findings establish KIF18A as a functional marker and mediator of mitotic instability in SCLC and reveal SAC activity as a key determinant of KIF18A inhibitor sensitivity.

## Materials and Methods

### Cell lines and culture

All cell lines were obtained from ATCC (https://www.atcc.org/), unless otherwise stated. NCI-H841, NCI-H1048 and DMS-53 were cultured in Dulbecco’s Modified Eagle Medium/Nutrient Mixture F-12 (DMEM/F-12), DMS273 was cultured in DMEM and other cell lines were cultured in Roswell Park Memorial Institute (RPMI)1640. The RA22#12 cell line was generated from tumor tissue collected during autopsy from a patient with metastatic small cell lung cancer. All media were supplemented with 10% fetal bovine serum (FBS). All cell lines were maintained at 37 °C with 5% CO₂ and were confirmed to be mycoplasma-negative prior to use.

### Generation of Cell lines expressing H2B-GFP and H2B-RFP

Lentiviruses were produced in HEK-293T cells by transient transfection with pCMV-VSV-G (8454, Addgene), pCMV-dR8.2 dvpr (8455, Addgene), and pHIV-H2B-eGFP (91776, Addgene) or pHIV-H2B-mRFP (18982, Addgene) using TurboFectin 8.0 (TF81001, OriGene). Viral supernatants were collected and used to infect target cells in the presence of polybrene (2 μg/mL). GFP- or RFP-positive populations were isolated by flow cytometry and confirmed by fluorescence microscopy.

### Inhibitors treatments

Cells were treated with the KIF18A inhibitor AM-9022 (synthesized by Amgen) at the indicated concentrations and time points. The MPS1 inhibitor reversine (656820-32-5, MedChemExpress) was used at 0.2 μM to induce chromosomal instability or at 0.05 μM for 48 h to reduce SAC activity. To activate the SAC, cells were treated overnight with 50 ng/ml nocodazole (HY-13520, MedChemExpress) or 10 mM monastrol (S8439, Selleckchem) for 3 hrs.

### Immunoblotting

Cells were lysed in RIPA buffer supplemented with protease and phosphatase inhibitor cocktails and PMSF. Protein concentrations were determined using a Bio-Rad protein assay. Equal amounts of protein were resolved by SDS–PAGE, transferred to PVDF membranes, and probed with primary antibodies against KIF18A (A301-080A, Bethyl Laboratories) and α-tubulin (T9026, MilliporeSigma) followed by HRP-conjugated secondary antibodies. Signals were detected by chemiluminescence, and band intensities were quantified using ImageJ software and normalized to α-tubulin.

### Immunostaining

Cells grown on glass coverslips were fixed with ice-cold methanol, washed with PBS, and blocked with 2% BSA in PBST. Cells were incubated with primary antibodies against Mad1 (MABE867, Sigma-Aldrich), BubR1 (EPR22544-48, Abcam), KIF18A (A301-080A, Bethyl Laboratories) and α-tubulin (T9026, MilliporeSigma) overnight at 4°C followed by fluorophore-conjugated secondary antibodies Alexa Fluor 488–conjugated anti-mouse IgG (A11029) and Alexa Fluor 568–conjugated anti-rabbit IgG (A11036; Invitrogen) . Nuclei were counterstained with DAPI, and coverslips were mounted using antifade mounting medium. Images were acquired by fluorescence microscopy were used as secondary antibodies.

### Microscopy and quantitative immunofluorescence Image analysis

For MAD1 and BubR1 signal intensity measurement, MAD1 and BubR1 immunostained cells were imaged on Delta Vision Core system (Applied Precision/GE Healthcare) consisting of an Olympus IX70 inverted microscope (Olympus America) with 100× NA 1.4 oil immersion objective and a CoolSnap HQ 12-bit camera (Photometrics) controlled by Softworx software. Z-stacks of at least 10 focal planes, each of 10μm were acquired. Signal intensity was measured using the plot profile tool in Softworx. To calculate fluorescence intensities, boxes of 8 × 8 pixels were drawn on kinetochore regions, as ascertained by co-localized foci of MAD1 and BubR1. For background, four boxes of 8 × 8 pixels were drawn at four random areas within the cytoplasm in the same cell. The maximum intensity values from all drawn areas were obtained using the data inspector tool in Softworx. Final fluorescence intensity for each protein was calculated by subtracting the average background intensity. For MAD1 signal intensity analysis in interphase cells, signal intensities were measured at five different spots in nuclear membrane in a cell using point tool in Fiji. Average intensity from 5 spots were calculated to obtain average MAD1 nuclear membrane signal intensities in a cell.

### Time-lapse imaging

For time-lapse imaging, cells were seeded in μ-Slide 8 well coverglass bottom chambers (ibidi) and treated with AM-9022 and/or reversin as experimental regime. Live-cell imaging was performed using a Nikon SoRa spinning disk confocal microscope equipped a 40x plan-apochromat objective lens (N.A. 0.95), Fusion BT sCMOS camera (Hamamatsu Photonics K.K.), and stage top incubator (Tokai Hit USA, Inc.) to maintain humidity, temperature at 37 °C and 5% CO₂. Z-stacked (0.16 um X-Y pixel size and 1.5 um z-step size) images were acquired from multiple positions in each well every 4 min for 48 h. Image analysis were performed using NIS-Elements software (v. 5.42; Nikon Instruments, Inc.).

### Cell viability assay (ATPlite)

Cell viability was assessed using the ATPlite luminescence assay (PerkinElmer) according to the manufacturer’s instructions. Cells were treated with serial dilutions of AM-9022 for 72 h and luminescence was measured using a microplate reader. IC□□ values were calculated by nonlinear regression using GraphPad Prism.

### Cell viability assay (MTS)

Cell viability was also determined using the MTS colorimetric assay (APExBIO). Following 72 h treatment with serial dilutions of AM-9022, absorbance at 490 nm was measured, IC□□ values were calculated using GraphPad Prism.

### Cell growth assay

Cell proliferation was monitored over 72 h following AM-9022 treatment using the Incucyte live-cell imaging system (Sartorius). Phase-contrast images were acquired every 6 h, and confluence was quantified automatically using the Incucyte software and plotted as a function of time.

### Doubling time and proliferation rate

Cell doubling times were measured following the manufacturer’s instructions (Selector Biosciences). Viable cells were counted using ViaStain™ AOPI Staining Solution (Revvity), replated at a defined density (Xb) and recounted after a defined interval (Xe). Doubling time (Td) was calculated as Td = (T × ln(2)) / ln(Xe / Xb).

### shRNA-mediated knockdown

KIF18A knockdown was achieved using lentiviral shRNA vectors (SVSHU6T16-L, Cellecta). Cells were transduced according to the manufacturer’s instructions, and stable knockdown cell lines were established by selection for RFP expression and puromycin resistance. shRNA expression was induced with doxycycline (8 μg/mL), and knockdown efficiency was confirmed by immunoblotting. shRNA sequences are listed in **Supplementary Table S1**.

### Bioinformatics and transcriptomic analysis

Public SCLC cell line expression data were obtained from the Cancer Cell Line Encyclopedia (CCLE) and SCLC CellMinerCDB. For RNA-seq generated in this study, total RNA was isolated from SCLC cell lines using the RNeasy kit (Qiagen, 74134) per the manufacturer’s protocol. RNA integrity was assessed on an Agilent 4150 TapeStation, and samples meeting quality thresholds were submitted to Novogene for library preparation and sequencing. Poly(A)-selected libraries were constructed and sequenced on an Illumina NovaSeq platform (paired-end 150 bp), yielding ∼6 Gb of raw data per sample. Raw FASTQ files underwent quality control using FastQC, followed by adapter trimming and quality filtering using Trimmomatic. Filtered reads were aligned to the human reference genome using STAR (v2.7.11b). Gene-level counts were quantified with featureCounts (Rsubread). Downstream analyses were performed in R (v4.2.3). Differential expression between AM-9022–sensitive and –resistant cell lines was performed using DESeq2, and genes with |log2 fold-change| ≥ 1.5 and nominal P < 0.05 were carried forward for pathway analyses (unless otherwise specified). Gene set enrichment analysis (GSEA) was performed using the Broad Institute Molecular Signatures Database (MSigDB) gene sets. Where indicated, single-sample GSEA (ssGSEA) was used to score predefined signatures (e.g., CES) across cell lines.

### Statistics

Statistical analyses of experimental data were performed using GraphPad Prism (version 10). Statistical tests were selected based on data distribution and sample size. For multiple-group comparisons, Kruskal–Wallis with Dunn’s or one-way ANOVA with Tukey’s tests were used as appropriate. For repeated measures, Friedman tests were applied. Two-way ANOVA was used for experiments involving two independent variables. Pairwise comparisons were performed using two-sided Wilcoxon rank-sum or Welch’s t-tests. Data are presented as mean ± SD unless otherwise indicated. A P value < 0.05 was considered statistically significant.

## Results

### KIF18A expression is associated with chromosomal instability and neuroendocrine features in SCLC

To investigate the genomic instability landscape of SCLC, we first compared indices that capture CIN and aneuploidy, including the fraction genome altered (FGA) (18), centromere and kinetochore gene Expression Score (CES) (19), and the CIN70 score (20), across lung cancer histological subtypes. Lung adenocarcinoma (LUAD) and lung squamous cell carcinoma (LUSC) primary tumor datasets were obtained from The Cancer Genome Atlas (TCGA) (21), and SCLC tumors were derived from the George et al. cohort (3).

SCLC tumors exhibited significantly higher FGA score compared with LUAD and LUSC (**Fig. 1A**). Consistently, SCLC tumors exhibited pronounced CIN, as evidenced by significantly increased CES and CIN70 scores, relative to LUAD and LUSC cohorts (**Fig. 1B**).

**Figure 1.**
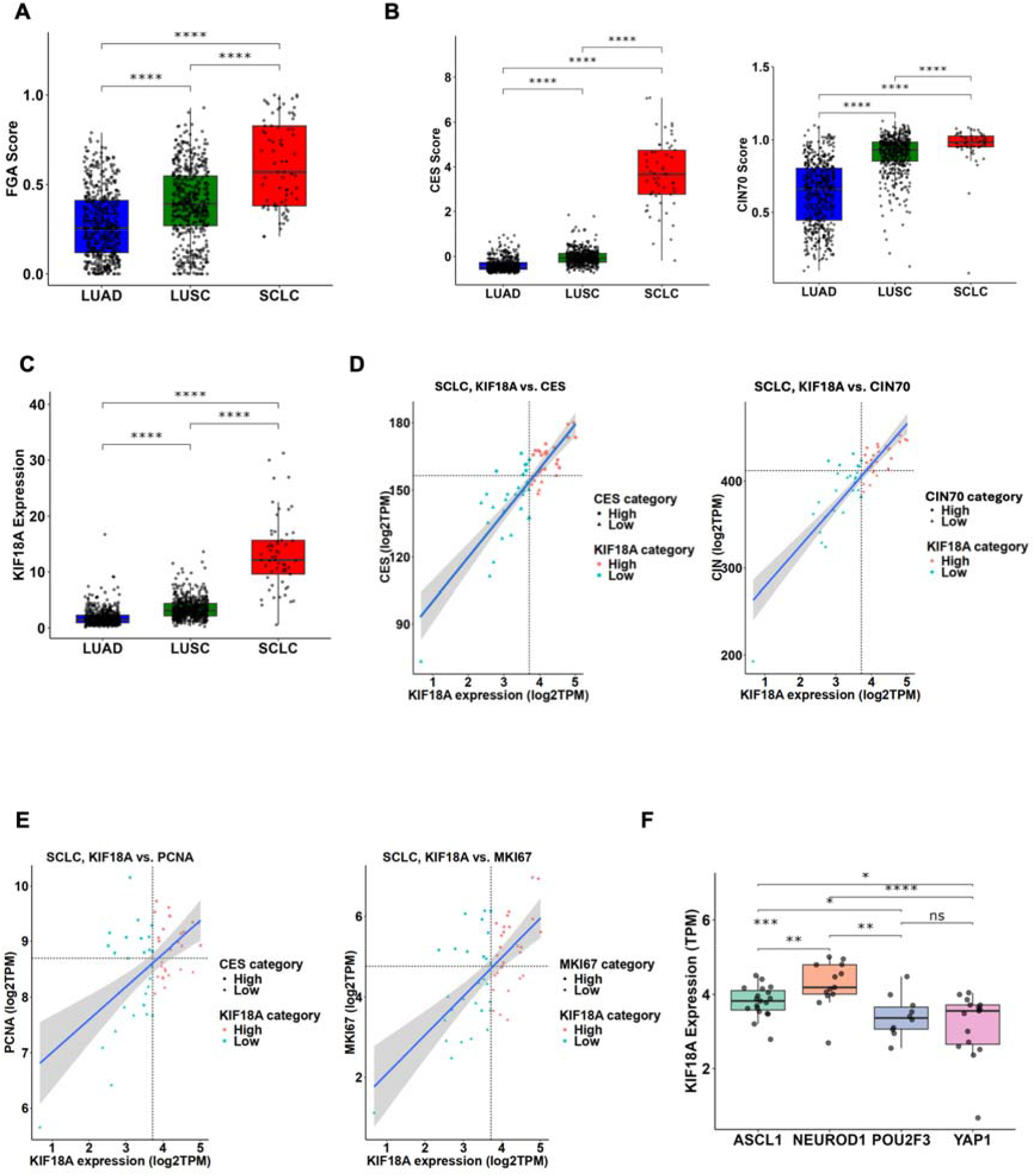
Association of KIF18A expression with chromosomal instability in SCLC. **A**, Box plot showing the fraction genome altered (FGA) scores across different lung cancer subtypes, including lung adenocarcinoma (LUAD, n=518) and lung squamous carcinoma (LUSC, n=500) from the TCGA database (21), and small cell lung carcinoma (SCLC, n=58) from George *et. al* (3). **B**, Box plots showing CES score; left (19) and CIN70 gene signature (right) (20) using TCGA (21) and George *et al.* datasets. **C**, Box plot showing KIF18A transcript levels across lung cancer subtypes in TCGA and George *et al*. datasets. **D**, Scatter plots showing the relationship between KIF18A transcript levels and CES (left) and CIN70 (right), across SCLC tumors from the George *et al*. dataset. **E**, Scatter plots showing correlations between KIF18A expression and proliferation markers PCNA (left) and MKI67 (right) in the George *et al*. dataset. **F**, Box plots showing KIF18A transcript levels across molecular subtypes of SCLC (ASCL1, NEUROD1, POU2F3, and YAP1) in the George et al. dataset. P values in panels A–C and F were calculated using two-sided Wilcoxon tests. Correlations in panels D and E were assessed using Spearman’s rank correlation. *, P < 0.05; **, P < 0.01; ***, P < 0.001; ****, P < 0.0001.

Given the pronounced CIN observed in SCLC, we next asked whether expression of the mitotic motor protein KIF18A, previously implicated as a selective dependency in genomically unstable cancers (16), was similarly enriched in SCLC and linked to CIN-related features. KIF18A expression was significantly upregulated in SCLC tumors compared to that with LUAD and LUSC (**Fig. 1C**). Within SCLC, KIF18A expression strongly correlated with CES (r = 0.83, P < 0.0001) and CIN70 (r = 0.85, P < 0.0001) scores, suggesting an association between KIF18A levels and the extent of CIN (**Fig. 1D**). We next investigated the relationship between KIF18A expression and proliferative capacity in SCLC. KIF18A expression also correlated positively with proliferation markers PCNA (r = 0.55, P < 0.0001) and MKI67 (r = 0.62, P < 0.0001) (**Fig. 1E**), linking KIF18A expression to proliferative drive in SCLC. Finally, analysis across SCLC molecular subtypes revealed differential expression, with KIF18A enriched in ASCL1-driven and NEUROD1-driven tumors, whereas POU2F3-driven and YAP-driven tumors expressed lower levels (**Fig. 1F**).

Together, these associations position KIF18A at the intersection of chromosomal instability, proliferative drive, and neuroendocrine identity in SCLC, raising the question of whether KIF18A expression marks a functional dependency or merely reflects an underlying CIN-high state. These tumor-level associations prompted us to examine whether KIF18A expression similarly varies across SCLC cell lines and whether it predicts functional dependency on KIF18A activity.

### SCLC cell lines exhibit heterogeneous KIF18A expression independent of drug sensitivity

KIF18A levels varied widely among SCLC cell lines (22). Consistent with tumor data (**Fig. 1C**), elevated KIF18A expression correlated with NE status (r = –0.28, P = 0.02; **Fig. 2A**). Varied mRNA expression levels of KIF18A across SCLC cell lines were validated experimentally. Quantitative PCR confirmed heterogeneous expression of KIF18A mRNA across representative cell lines (**Fig. 2B**). Notably, DMS53 cells expressed comparative levels of KIF18A as the control cell line, CRL9609. Consistent with transcript levels, Western blotting revealed variable protein levels of KIF18A across the panel; however, the differences in expression levels were not statistically significant (**Fig. 2C**).

**Figure 2.**
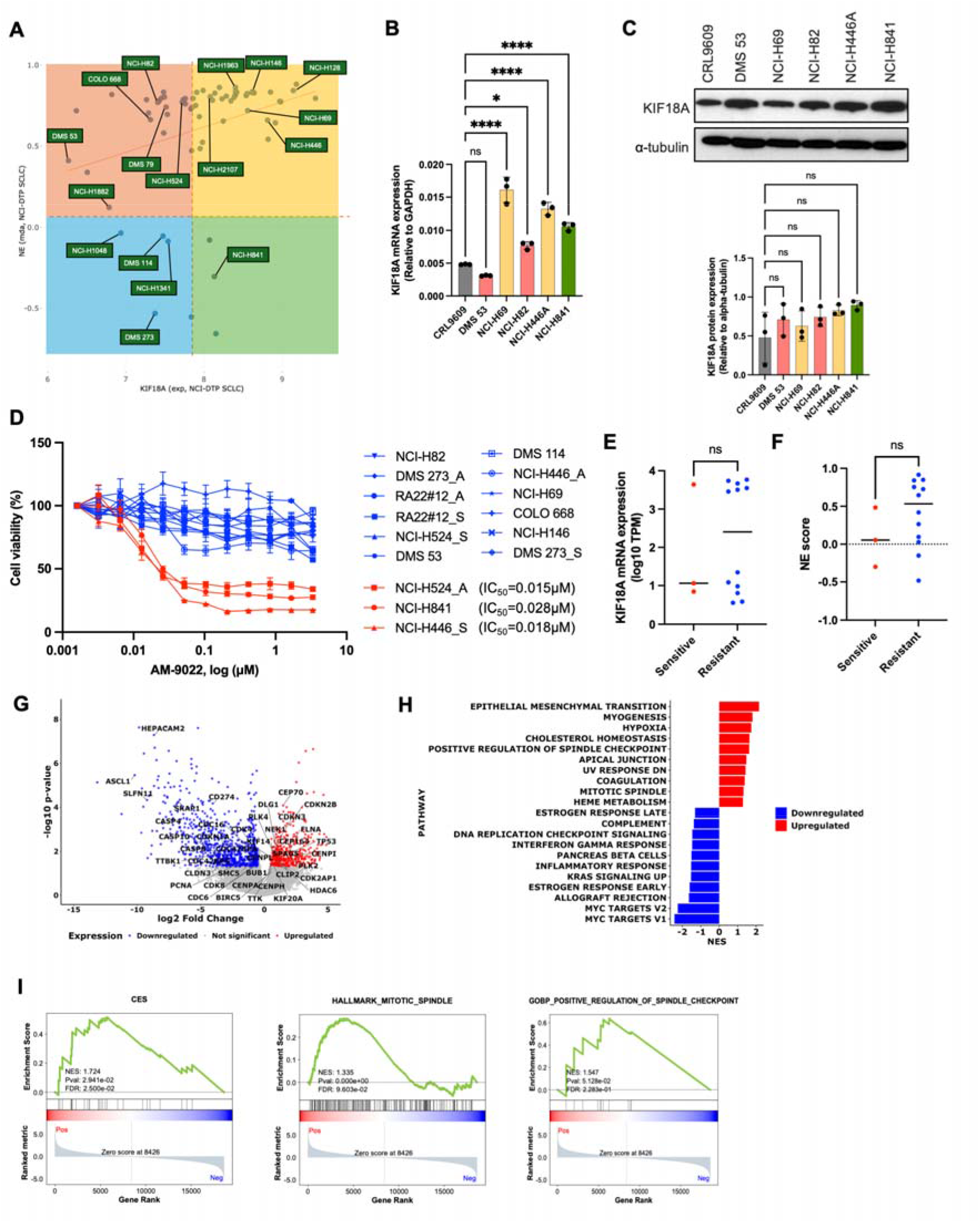
Molecular features associated with sensitivity to KIF18A inhibition in SCLC. **A**, Scatter plot showing the correlation between KIF18A mRNA expression and neuroendocrine (NE) status across SCLC cell lines from the CellMinerCDB SCLC dataset. Each datapoint represents an individual SCLC cell line. **B**, Bar graph showing relative KIF18A mRNA levels across SCLC cell lines as measured by quantitative RT-PCR. Expression was normalized to GAPDH. Data represent mean ± SD from three independent biological replicates. **C**, Western blot analysis of KIF18A protein levels across a panel of SCLC cell lines. α-Tubulin was used as a loading control. Bar graph showing relative KIF18A protein levels across SCLC cell lines as measured by immunoblotting. Protein expression was normalized to α-tubulin. Data represent mean ± SD from three independent biological replicates. **D**, Dose–response curves showing the sensitivity of SCLC cell lines to a KIF18A inhibitor, generated using technical replicates. IC_50_ values were used to classify cell lines as sensitive (<0.03 µM) or resistant (>3 µM). **E**, Dot plots showing KIF18A transcript levels for individual SCLC cell lines classified as KIF18A inhibitor–sensitive or –resistant. **F**, Dot plots showing neuroendocrine (NE) scores for individual SCLC cell lines classified as KIF18A inhibitor–sensitive or –resistant. **G**, Volcano plot showing differential gene expression between AM-9022–sensitive (n=3) and –resistant (n=17) SCLC cell lines based on RNA-seq analysis. Genes upregulated in sensitive cell lines are shown in red, whereas genes downregulated in sensitive cell lines (correspondingly upregulated in resistant cell lines) are shown in blue. Labeled genes indicate candidates associated with drug sensitivity or resistance. **H**, Summary of gene set enrichment analysis (GSEA) comparing transcriptomic profiles of AM-9022–sensitive (n=3) versus resistant (n=17) SCLC cell lines. Gene sets with positive normalized enrichment scores (NES), shown in red, are enriched in AM-9022–sensitive SCLC cell lines, whereas gene sets with negative NES, shown in blue, are enriched in resistant cell lines and depleted in sensitive cell lines. Sensitive lines display enrichment of SAC-related mitotic pathways. **I**, GSEA enrichment score plots showing enrichment of chromosomal instability (CES), mitotic spindle, and spindle checkpoint in AM-9022–sensitive SCLC cell lines relative to resistant cell lines. P values in panels B and C were calculated using one-way ANOVA with Dunnett’s multiple-comparison test. P values in panels E and F were calculated using two-sided Wilcoxon tests. *, P < 0.05; ****, P < 0.0001.

Since KIF18A inhibitor has been reported to be lethal to chromosomally unstable cells (16,23,24), next we tested if KIF18A inhibitor was uniformly lethal across SCLC cell lines with varying KIF18A expression. We analyzed a panel of 15 SCLC cell lines spanning a broad range of molecular and phenotypic features, including NE, non-NE, adherent or suspension growth patterns (**Supplementary Table 2**). Cells were treated with AM-9022, a selective ATP-competitive inhibitor of KIF18A that blocks its motor activity and impairs chromosome alignment during mitosis (23). Importantly, this mechanism is directly relevant to clinically advanced KIF18A inhibitors, including VLS-1488 and sovilnesib, which are currently undergoing Phase I clinical evaluation in patients with solid tumors (ClinicalTrials.gov: NCT05902988, NCT06084416) (25,26).

Dose–response analyses following 72-hour treatment revealed marked heterogeneity in sensitivity. Three cell lines (H841, H524A and H446S) exhibited low IC50 values and were classified as sensitive, whereas the majority of lines were resistant even at higher drug concentrations (**Fig. 2D**; **Supplementary Fig. S1A**). Importantly, neither KIF18A mRNA expression nor NE score differed significantly between sensitive and resistant cell lines (**Fig. 2E, F**). Consistent with this, neither showed a significant correlation with normalized AUC (KIF18A mRNA: r = 0.20, P = 0.46; NE score: r = 0.37, P = 0.16; **Supplementary Fig. S1B, C**). Similarly, KIF18A protein levels did not correlate significantly with normalized AUC (r = -0.77, P = 0.11; **Supplementary Fig. S1D**). Together, these results indicate that although SCLC cell lines exhibit substantial variability in KIF18A expression which tracks with NE status (**Fig. 1F**), neither baseline KIF18A expression nor NE score predicts sensitivity to KIF18A inhibition.

Given the essential role of KIF18A motor activity in chromosome alignment and mitotic progression, we hypothesized that sensitivity to KIF18A inhibition might reflect differences in cell-cycle kinetics that limit the accumulation of mitotic defects within the treatment window. To test this, we compared doubling time and proliferation rates between sensitive and resistant cell lines (**Supplementary Table 2**). No significant association was observed between doubling time and drugs sensitivity (r = 0.37, P = 0.45; **Supplementary Fig. S1E, Supplementary Table 3**), suggesting that factors beyond proliferation rate or KIF18A expression govern KIF18A dependency in SCLC.

To investigate the transcriptional programs that might underlie the observed heterogeneity in sensitivity to KIF18A inhibition, we performed RNA sequencing and compared the transcriptomes of AM-9022–sensitive and –resistant SCLC cell lines. Differential expression analysis identified numerous genes with significant expression changes between the two groups. Given that KIF18A is a microtubule-based motor protein essential for chromosome alignment and mitotic progression, we specifically focused on differentially expressed genes (DEGs) associated with cell cycle, SAC and microtubule dynamics (**Fig. 2G**). Sensitive cell lines exhibited significant upregulation of genes associated with centrosome maturation, mitotic spindle organization, and cell cycle regulation, including CEP70, PLK4, PLK2, NEK1, CENPI, and CDKN3. Genes associated with chromosomes movement (HEPACAM2, CDC42BPG), DNA damage signaling (SLFN11, CD274), and microtubule dynamics (TTBK1) were downregulated in these cell lines.

Pathway enrichment analysis further revealed significant enrichment of mitotic spindle regulation, checkpoint-associated cell cycle processes, and stress response–related programs in sensitive cell lines compared with resistant counterparts (**Fig. 2H**). Transcriptomic comparisons between AM-9022–sensitive and –resistant SCLC cell lines similarly demonstrated enrichment of genes involved in mitotic spindle organization and SAC regulation in the sensitive group (**Fig. 2I**). Notably, CIN–associated signatures, including the CES, were among the top-ranked pathways. In line with this, ssGSEA analysis showed that sensitive cell lines exhibited significantly higher CES scores than resistant ones (P = 0.01; **Supplementary Fig. S1F**).

Collectively, these analyses demonstrates that KIF18A inhibitor sensitivity is accompanied by a broad rewiring of mitotic spindle, checkpoint-related cell cycle processes and stress response–related programs. The prominence of spindle assembly checkpoint–associated gene sets in sensitive cell lines prompted further functional interrogation of mitotic checkpoint activity as a determinant of AM-9022 response.

### KIF18A inhibition selectively suppresses proliferation through defects in mitotic progression in sensitive SCLC cells

To functionally characterize the cellular response to KIF18A inhibition, we treated the AM-9022- sensitive H841 and resistant DMS53 SCLC cell lines with increasing concentrations of AM-9022, and assessed their proliferation, cell viability, and mitotic dynamics across multiple time points.

Incucyte based proliferation assays revealed a pronounced, dose-dependent suppression of proliferation in H841 cells, whereas DMS53 cells exhibited only marginal growth inhibition even at the highest tested concentrations (**Fig. 3A**). At 72 h, treatment with 0.04 µM AM-9022 reduced confluence in H841 cells from 74% to 46% (∼38% inhibition, P = 0.01), with no effect in DMS53 cells (P = 0.26). This differential response was corroborated by MTS cell viability assays, which showed a marked decrease in cell viability in H841 cells, with an IC□□ of 0.032 µM, in contrast to the largely unchanged viability profile observed in DMS53 cells (**Fig. 3B**). Collectively, these findings indicate that AM-9022 selectively impairs proliferative capacity in sensitive SCLC cells.

**Figure 3.**
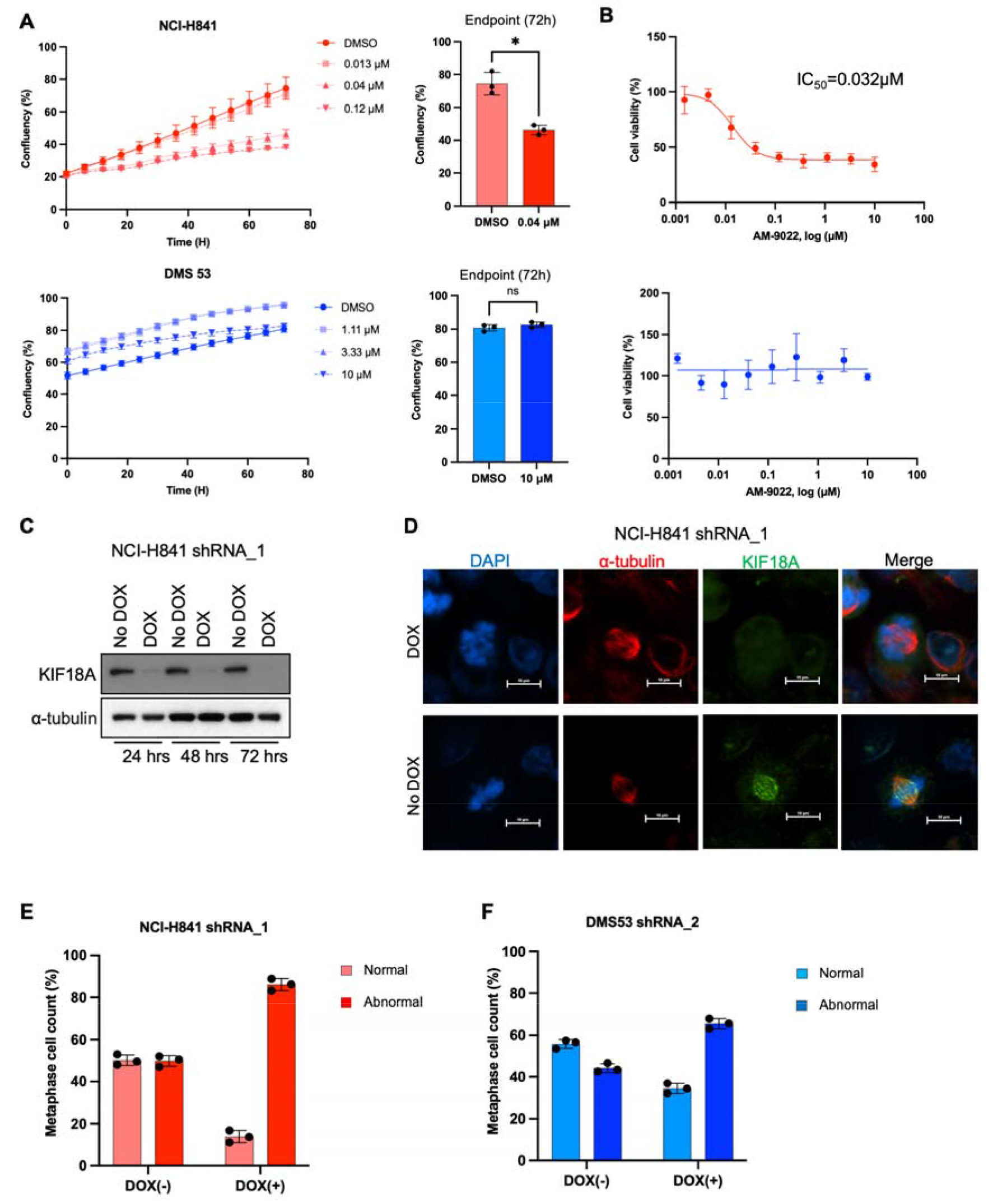
KIF18A inhibition selectively impairs proliferation and mitotic spindle integrity in sensitive SCLC cells. **A**, Incucyte live-cell imaging analysis for cell proliferation in NCI-H841 (upper) and DMS53 (lower) SCLC cell lines treated with different concentrations of AM-9022 for multiple time points. Dot plots displaying endpoint confluency at 72 h in NCI-H841 (upper) and DMS53 (lower) cells treated with indicated concentrations of AM-9022. P values were calculated using Welch’s test. *, P < 0.05. **B**, Cell viability measured by MTS assay following AM-9022 treatment in NCI-H841 and DMS53 cells. H841 cells show a marked reduction in viability, whereas DMS53 cells remain relatively resistant. IC_50_ values derived from dose–response curves are indicated (H841 ≈ 0.032 μM; DMS53 > 10 μM). **C**, Immunoblot analysis of KIF18A protein levels following doxycycline (DOX)–induced shRNA expression in NCI-H841 cells. Cells were cultured in the absence or presence of DOX (8 μg/mL) for the indicated times (24, 48, and 72 h). α-Tubulin was used as a loading control. **D**, Representative immunofluorescence images of mitotic NCI-H841 cells following DOX-induced KIF18A knockdown using shRNA_1. Cells were immunostained with α-tubulin (red), and KIF18A (green). DNA was stained with DAPI (blue). Scale bars, 10 μm. **E**, Quantification of metaphase cells with normal or abnormal spindle organization following KIF18A knockdown in NCI-H841 cells. For each biological replicate, 100 metaphase cells were scored and classified as normal or abnormal. The percentage of cells in each category is shown. Bars indicate dots represent individual biological replicates (n = 3). **F**, Quantification of metaphase cells with normal or abnormal spindle organization following KIF18A knockdown in DMS53 cells. For each biological replicate, 100 metaphase cells were scored and classified as normal or abnormal. The percentage of cells in each category is shown. Bars indicate the mean, and dots represent individual biological replicates (n = 3).

To delineate the mechanistic basis of this differential sensitivity, we examined mitotic progression following KIF18A inhibition, building on prior reports of spindle and chromosome congression defects upon KIF18A loss (13). Mitotic index was quantified in two KIF18A sensitive and two KIF18A resistant cell lines following treatment with AM-9022 for 72 h. KIF18A inhibition resulted in a markedly increased mitotic index in sensitive cell lines, indicative of mitotic arrest (**Supplementary Fig. S2A**). In contrast, resistant cell lines exhibited a low mitotic index upon KIF18A inhibition (**Supplementary Fig. S2A**). These finding indicate that resistant cells fail to undergo mitotic arrest in response to KIF18A inhibition.

To determine whether these mitotic defects reflect an on-target requirement for KIF18A, we next examined the consequences of genetic depletion of KIF18A. We generated H841 and DMS53 cells harboring a doxycycline-inducible knockdown of KIF18A and optimized the concentration and duration of doxycycline treatment, identifying 8 μg/mL as the condition that achieved efficient depletion (**Supplementary Fig. S2B and S2C**). Robust knockdown of KIF18A was observed following doxycycline treatment, as confirmed by immunoblotting (**Fig. 3C**) and immunofluorescence analyses (**Fig. 3D**). Genetic depletion of KIF18A resulted in a significant increase in the proportion of abnormal metaphase cells (**Fig. 3E**). In contrast, KIF18A depletion in resistant cell lines led to a more modest increase in abnormal metaphase cells (**Fig. 3F**). Collectively, these results demonstrate that KIF18A is required for proper metaphase chromosome alignment and mitotic progression in sensitive SCLC cells.

### KIF18A inhibition induces SAC-dependent mitotic arrest and cell death in sensitive SCLC cell line

Having established that KIF18A is required for proper metaphase chromosome alignment, we next used time-lapse imaging to examine how KIF18A inhibition affects mitotic progression and cell fate in sensitive SCLC cell lines. H841 cells stably expressing RFP-tagged histone H2B (H841-H2B-RFP) were treated with AM-9022 for 8 h prior to imaging, followed by continuous time-lapse acquisition for 48 h (**Fig. 4A**). Fluorescently tagged H2B was used as a nuclear marker to assess chromosome dynamics and segregation fidelity.

**Figure 4.**
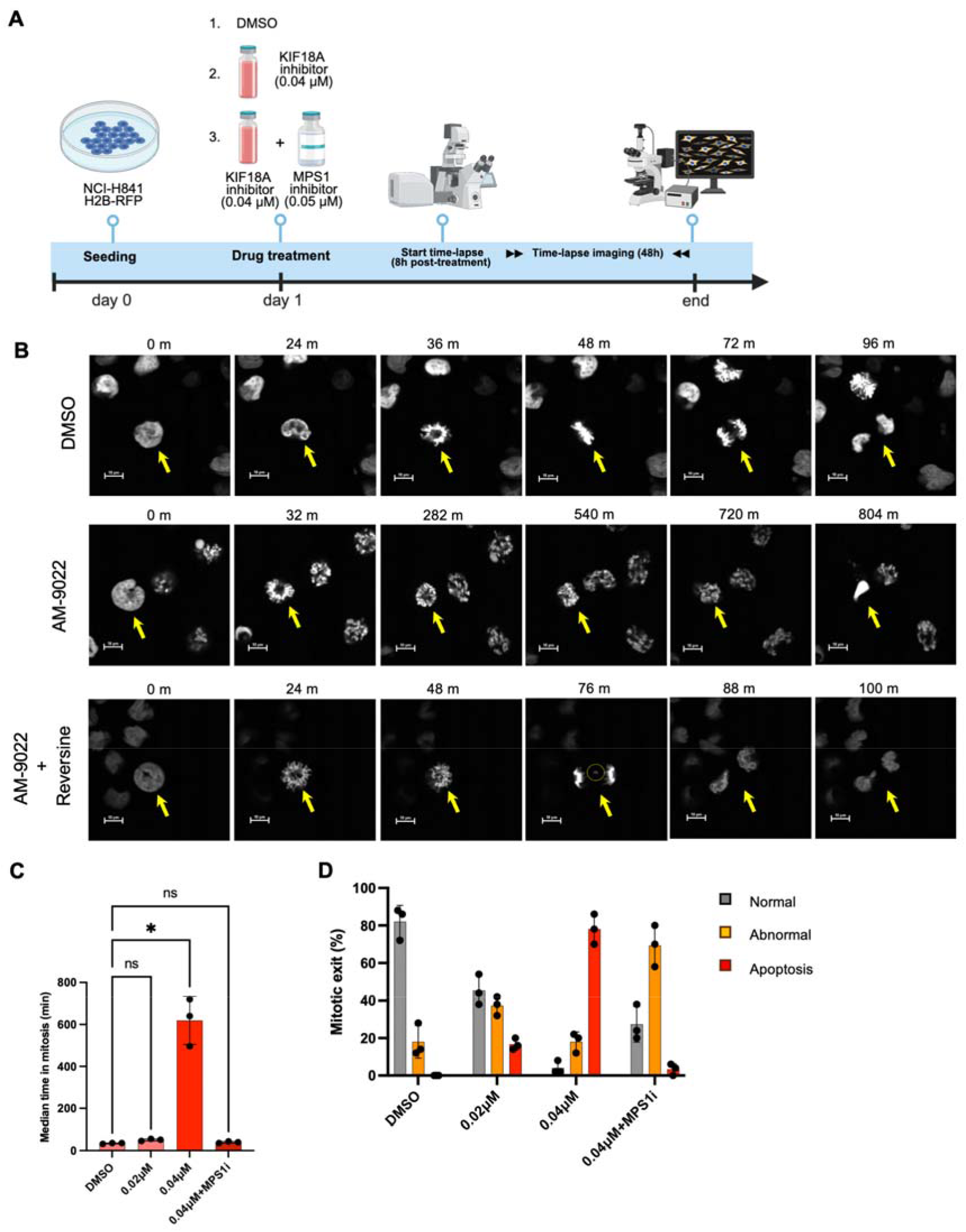
SAC-dependent mitotic arrest and cell fate outcomes following KIF18A inhibition in sensitive SCLC cells. **A**, Schematic timeline illustrating the experimental workflow for time-lapse imaging. **B**, Representative time-lapse images of H2B-RFP expressing NCI-H841 cells treated with DMSO, AM-9022 (0.04 μM), or AM-9022 in combination with the SAC inhibitor reversine (0.05 μM) showing mitotic outcome. Yellow arrows indicate mitotic cells. Scale bars, 10 μm. **C**, Quantification of mitotic timing in NCI-H841 cells following the indicated treatments. Mitotic timing was calculated by counting frames, each frame equivalent to 4 minutes, from NEBD to anaphase onset or cell death in time-lapse images. Bars represent the median, and dots indicate individual biological replicates (n=3). P values were calculated using Friedman test with Dunn’s multiple-comparison test. *, P < 0.05. **D**, Quantification of cell fate outcomes in NCI-H841 cells following the indicated treatments. Bar graphs show the percentage of cells undergoing normal or abnormal mitotic exit, or apoptosis. Dots indicate individual biological replicates (n = 3).

Representative time-lapse images illustrate mitotic timing from nuclear envelope breakdown (NEBD) until either entry into anaphase or cell death across the indicated treatment conditions (**Fig. 4B**). Upon treatment with AM-9022, cells exhibited prolonged mitotic arrest, ultimately culminating in apoptotic cell death. Quantitative analysis demonstrated that AM-9022 induced a dose-dependent mitotic delay in H841-H2B-RFP cells (**Fig. 4C**), followed by either mitotic slippage with aberrant chromosome segregation or cell death, predominating at higher drug concentrations (**Fig. 4D, Supplementary Fig. S3**). In contrast, most DMSO-treated cells progressed through mitosis normally.

To test whether this prolonged mitotic arrest was dependent on SAC signaling, cells were co-treated with a low dose of reversine (0.05 mM), an MPS1 inhibitor and AM-9022. **4C).** Notably, co-treatment, which partially overrides SAC signaling, restored mitotic timing comparable to DMSO control **(Fig. 4C).** However, under these conditions, a substantial fraction of cells exited mitosis with abnormal chromosome segregation **(Fig. 4D)**, indicating that partial SAC override permits mitotic exit despite persistent segregation defects in AM-9022-treated H841 cells.

Collectively, these findings indicate that KIF18A inhibition triggers sustained SAC activation, leading to prolonged mitotic arrest and cell death in AM-9022 sensitive SCLC cells. Pharmacologic attenuation of SAC signaling abrogates mitotic arrest, underscoring SAC activation as a key mediator of KIF18A inhibitor–induced lethality.

### Reduced MAD1/BubR1 recruitment and impaired SAC activation in resistant cells

We next investigated whether the defects in mitotic arrest upon treatment with AM-9022 in resistant DMS53 cells reflects intrinsic defects in SAC signaling. To investigate this, DMS53 and H841 cells were treated with monastrol, an EG5 (kinesin-5) inhibitor to enrich the population prometaphase cells, a point at which SAC activation peaks due to the presence of tensionless kinetochores (27).

Cells were immunostained with MAD1 and BubR1, core components of the mitotic checkpoint complex, followed by localization and quantitative analysis of MAD1 and BubR1 at prometaphase kinetochores. As expected for SAC-active state, MAD1 and BubR1 localizations at kinetochores were intact in H841 cells, indicating intact SAC signaling. In contrast, DMS53 cells exhibited reduced recruitment of MAD1 and BubR1 at kinetochores, suggesting compromised SAC function (**Fig. 5A, B**). Quantitative immunofluorescence analysis confirmed that the signal intensities of MAD1 and BubR1 at prometaphase kinetochores were significantly reduced in DMS53 cells compared to H841 cells (**Fig. 5C, D**), demonstrating a deficient SAC signaling in resistant cells. These findings were reproducible across additional SCLC models, with similar results observed in H524A (sensitive) and DMS114 (resistant) cell lines **(Supplementary Fig. S4A-D).** Importantly, MAD1 signal intensities at nuclear membrane in DMS53 and H841 cell were comparable **(Fig. S4E, F)**, confirming that not overall, but only the kinetochore recruitment of MAD1 was affected in resistant cells.

**Figure 5.**
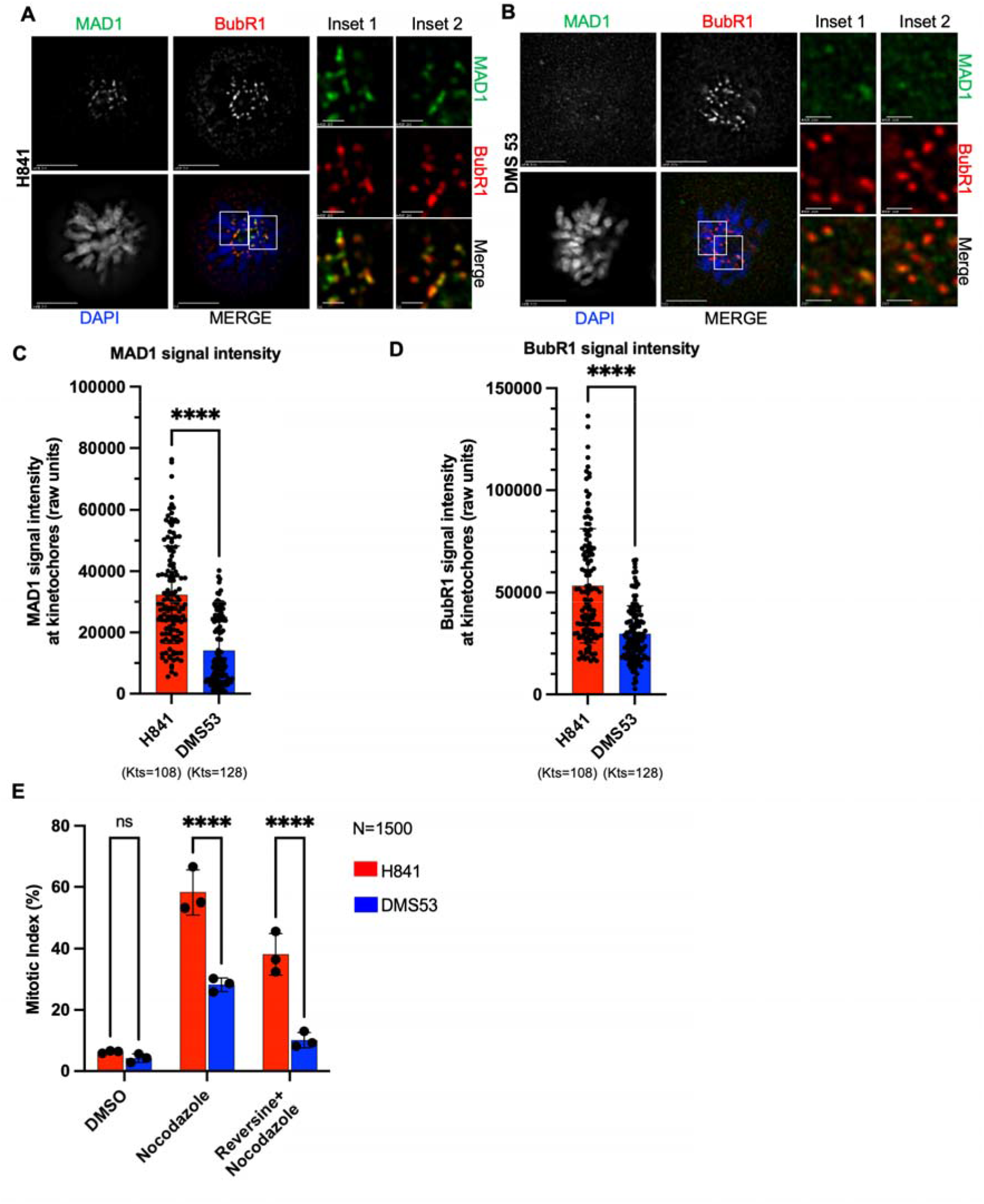
Differential spindle assembly checkpoint signaling at kinetochores in KIF18A inhibitor–sensitive and –resistant SCLC cells. **A**, Representative immunofluorescence images showing localization MAD1 (green) and BubR1 (red) at prometaphase kinetochores in monastrol treated NCI-H841 cells. Insets show zoomed images from white boxed areas in main images. Scale bar: 5 μm for main images and 1 μm for insets. **B**, Representative immunofluorescence images showing localization of MAD1 (green) and BubR1 (red) localization at prometaphase kinetochores in monastrol treated DMS53 cells. Insets show zoomed images from white boxed areas in main images. Scale bar: 5 μm for main images and 1 μm for insets. **C**, Quantification of prometaphase kinetochore-associated MAD1 signal intensities in monastrol treated NCI-H841 and DMS53 cells. Bar represents mean ± SD across kinetochores (n=128 for H841 and n=108 for DMS53). Kinetochores from at least 12 cells from two independent experiments were analyzed. Each dot represents a single kinetochore. **D**, Quantification of prometaphase kinetochore-associated BubR1 signal intensities in monastrol treated NCI-H841 and DMS53 cells. Bar represents mean ± SD across kinetochores (n=128 for H841 and n=108 for DMS53). Kinetochores from at least 12 cells from two independent experiments were analyzed. Each dot represents a single kinetochore. **E**, Quantification of mitotic index following nocodazole treatment in NCI-H841 and DMS53 cells. Cells were treated with DMSO, nocodazole, or nocodazole in combination with the SAC inhibitor reversine. 1500 cells were counted for each cell line and for each condition across three biological replicates. Dots indicate individual biological replicates (n = 3). Data are presented as mean ± SD. P values were calculated using two-way ANOVA with cell line and treatment as factors. P values in panels C and D were calculated using Welch’s test. ****, P < 0.0001.

To functionally validate these observations, we challenged cells with a low dosage of nocodazole to induce microtubule depolymerization. In SAC-proficient cells, nocodazole treatment enforces mitotic arrest, whereas SAC-deficient cells undergo premature mitotic exit. Consistent with intact checkpoint function, H841 cells displayed a pronounced mitotic arrest. In contrast, DMS53 cells failed to sustain mitotic arrest under the same conditions (**Fig. 5E)**. Notably, DMS53 cells treated with nocodazole alone exited mitosis prematurely than H841 cells co-treated with nocodazole and reversine, indicating that SAC in DMS53 cells is intrinsically inefficient, even relative to pharmacologically induced SAC-deficient H841 cells. Together, these data demonstrate that DMS53 cells are functionally deficient in SAC activation, both at the level of kinetochore signaling and checkpoint response, providing a mechanistic explanation for their resistance to KIF18A inhibition–induced mitotic arrest. Collectively, these results establish impaired SAC signaling as a reproducible feature of KIF18A inhibitor–resistant SCLC cell lines.

### Induction of CIN partially sensitizes resistant SCLC cells to KIF18A inhibition

Although resistant cells exhibit attenuated SAC signaling (**Fig. 5**), we reasoned that further increasing CIN might exceed the tolerance threshold and expose a dependence on mitotic fidelity mechanisms, including KIF18A. We therefore DMS53 cells with a low dose of reversine to induce CIN and evaluated whether this manipulation alters their response to KIF18A inhibition.

We employed a sequential treatment strategy in DMS53 cells (**Fig. 6A**). Cells were first pre-treated with reversine to induce CIN. Following drug washout and recovery for 24 hrs, cells were subsequently treated with AM-9022, and long-term live-cell time-lapse imaging was performed. Representative time-lapse images illustrate mitotic behavior under this sequential treatment condition (**Fig. 6B**).

**Figure 6.**
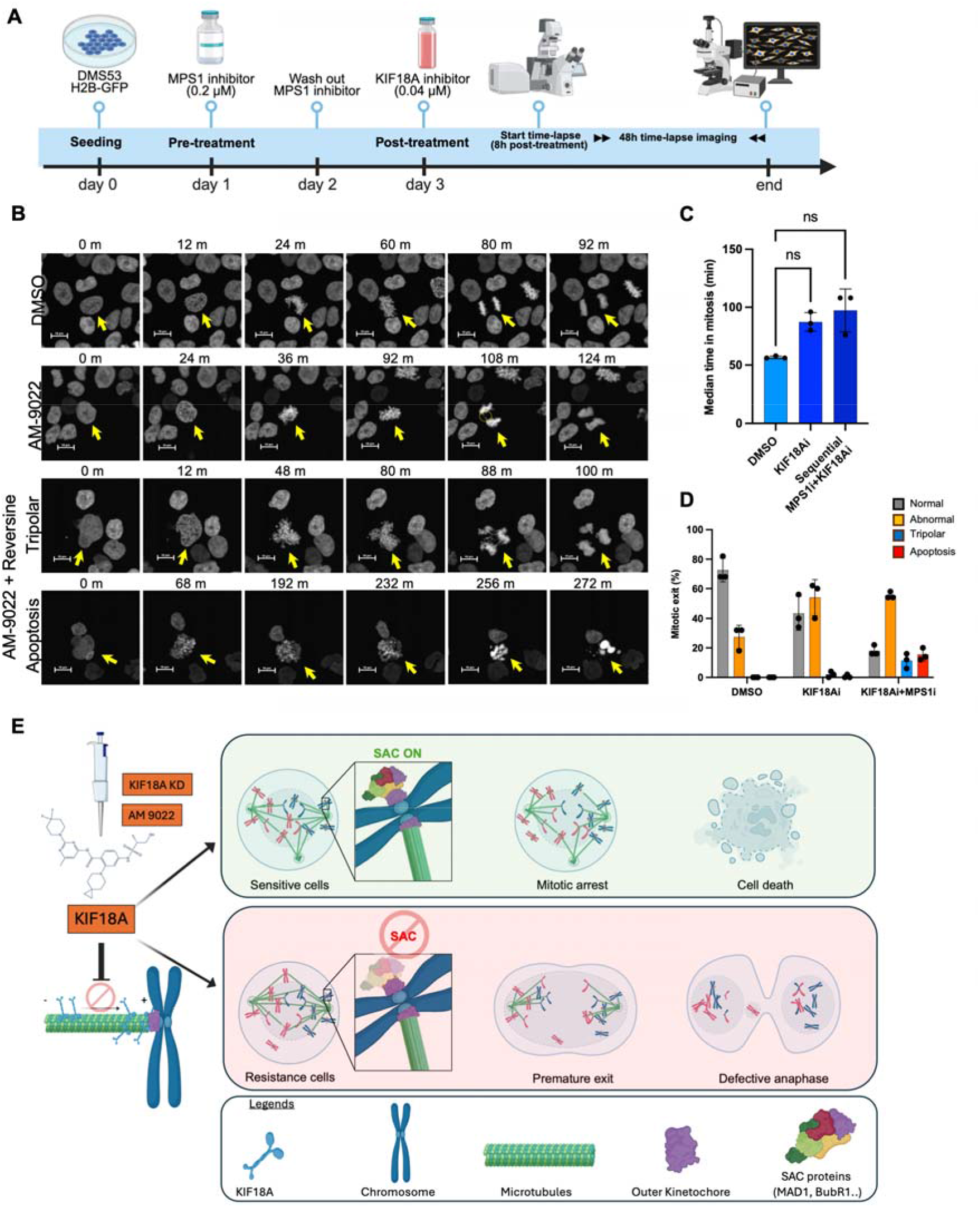
SAC integrity determines cellular responses to KIF18A inhibition in SCLC. **A**, Schematic timeline illustrating the experimental workflow for time-lapse imaging. **B**, Representative time-lapse images of H2B-GFP expressing DMS53 cells treated with DMSO, AM-9022 (0.04 μM), or AM-9022 following pre-treatment with the SAC inhibitor reversine. Representative frames from the time-lapse sequences are shown. Yellow arrows indicate mitotic cells. Scale bars, 10 μm. **C**, Quantification of mitotic timing in DMS53 cells following the indicated treatments. Mitotic timing was calculated by counting frames, each frame equivalent to 4 minutes, from NEBD to anaphase onset or cell death in time-lapse images. Bars represent the median, and dots indicate individual biological replicates (n=3). P values were calculated using Friedman test with Dunn’s multiple-comparison test. **D**, Quantification of cell fate outcomes in DMS53 cells following the indicated treatments. Bar graphs show the percentage of cells undergoing normal or abnormal mitotic exit, tripolar, or apoptosis. Dots represent individual biological replicates (n = 3). **E**, Proposed model depicting how SAC integrity shapes cellular responses to KIF18A inhibition in SCLC. In AM-9022–sensitive cells, KIF18A inhibition causes chromosome misalignment, prolonged SAC activation, mitotic arrest, and cell death. In contrast, resistant cells exhibit attenuated SAC signaling and are able to progress through mitosis despite KIF18A inhibition, allowing continued survival and proliferation.

In the resistant DMS53 cell line, treatment with AM-9022 alone did not result in a significant prolongation of mitotic timing, and sequential pre-treatment with reversine led to only a modest, non-significant extension of mitosis (**Fig. 6C**). However, quantitative analyses revealed that a subset of DMS53 cells underwent cell death following reversine pre-treatment and subsequent AM-9022 treatment, whereas AM-9022 treatment alone failed to induce detectable cell death in this model (**Fig. 6D, Supplementary Figure S5**). Together, these results indicate that induction of CIN partially re-sensitizes resistant SCLC cells to KIF18A inhibition, supporting a checkpoint-dependent, inducible vulnerability to mitotic perturbation.

## Discussion

SCLC is one of the most chromosomally unstable human malignancies, characterized by pervasive aneuploidy, high replication stress, and rapid proliferative turnover (3,18). Such extreme CIN has been proposed to generate targetable mitotic vulnerabilities (16). Among these, the mitotic kinesin KIF18A, a kinesin-8 family motor that suppresses microtubule plus-end dynamics and promotes chromosome alignment, has emerged as a selective dependency in CIN-high tumors (23). In this study, we investigated whether this CIN-associated dependency extends to SCLC, a prototypical CIN-high cancer.

We found that among lung cancer histological subtypes, KIF18A expression was particularly elevated in SCLC, and strongly correlated with chromosomal instability-associated gene signatures such as CES (19), CIN70 (20), as well as and proliferation markers (MKI67, PCNA). This pattern suggests that KIF18A upregulation accompanies the elevated mitotic activity and genomic plasticity that typify SCLC, particularly in ASCL1- and NEUROD1-driven neuroendocrine subtypes (11).

However, despite this correlation, KIF18A expression alone did not predict dependency. Across a panel of SCLC cell lines, differences in KIF18A mRNA and protein expression were observed; however, these differences did not correlate with sensitivity to the selective inhibitor AM-9022, indicating that dependency is determined not by expression abundance but by the mitotic context in which KIF18A functions. This observation extends prior studies by revealing a mechanistic link between spindle checkpoint competence and KIF18A dependency. While previous reports demonstrated that chromosomally unstable cancers rely on KIF18A for proliferation (16) and that perturbation of spindle checkpoint signaling through TTK inhibition can sensitize mitotically aberrant tumors to therapy (28), our findings provide the first evidence that the functional status of the spindle assembly checkpoint dictates sensitivity to KIF18A inhibition in CIN-high SCLC.

Transcriptomic profiling between AM-9022–sensitive and –resistant cell lines revealed distinct transcriptional programs involving mitotic spindle organization and checkpoint regulation. Sensitive lines displayed enrichment of hallmark pathways related to chromosomal instability, spindle assembly, and cell-cycle progression, together with higher CES scores. In contrast, resistant lines exhibited reduced expression and kinetochore localization of core SAC components MAD1 and BUBR1, consistent with defective checkpoint signaling. Such phenotypes resemble those in SAC-compromised models, where cells prematurely exit mitosis despite spindle perturbation (13). Accordingly, live-cell imaging demonstrated that KIF18A inhibition triggered pronounced mitotic arrest and apoptosis in SAC-proficient cells, whereas SAC-deficient cells underwent only mild mitotic delay and survived despite chromosome segregation errors. Together, these findings establish SAC competence as a critical determinant of vulnerability to KIF18A inhibition.

Mechanistically, these results support a model in which SAC-proficient SCLC cells rely on KIF18A to maintain spindle tension and alignment fidelity under conditions of chromosomal instability. When KIF18A is inhibited, persistent kinetochore–microtubule attachment defects activate SAC signaling and trigger apoptotic death, whereas SAC-deficient cells fail to sustain arrest and instead proceed through erroneous mitosis, preserving viability (**Fig. 6E**). This is consistent with recent work showing that SAC defects confer tolerance to mitotic stressors such as spindle poisons (29,30).

From a translational perspective, SAC integrity may serve as a functional biomarker for predicting response to KIF18A-targeted therapy. Given the heterogeneous SAC status across SCLC subtypes, therapeutic combinations that modulate checkpoint signaling could enhance efficacy. Indeed, our data support a synthetic-lethal framework in which transient amplification of chromosomal instability, achieved by sequential reversine treatment followed by AM-9022, can resensitize resistant cells by exceeding their tolerance for mitotic errors. Such “CIN-escalation” approaches may extend the therapeutic reach of KIF18A inhibition beyond SAC-proficient tumors, aligning with emerging concepts that controlled CIN elevation can potentiate mitotic stress therapies (31).

Several limitations should be acknowledged. First, the number of cell lines analyzed, although representative, remains limited for capturing the full molecular diversity of SCLC. Second, mechanistic findings were derived from in vitro models; in vivo validation using PDX or organoid systems will be essential to confirm SAC proficiency as a predictive biomarker. Third, AM-9022 remains a preclinical compound with incompletely defined pharmacologic and off-target profiles. Finally, while SAC function emerged as the dominant determinant of response, emerging evidence suggests that other mitotic regulators or metabolic adaptations may also modulate KIF18A dependency (32). Future work integrating proteogenomic and phospho-signaling analyses in clinical SCLC samples will be crucial to refine predictive models for KIF18A-targeted therapy.

In summary, this study delineates a mechanistic link between chromosomal instability, SAC competence, and KIF18A dependency in SCLC. KIF18A inhibition selectively induces mitotic arrest and apoptosis in SAC-proficient, CIN-high cells, whereas SAC-deficient cells bypass arrest and continue proliferating despite aberrant segregation. These findings position SAC integrity as both a biomarker and a therapeutic lever in SCLC and provide a rational basis for KIF18A-directed strategies in genomically unstable yet checkpoint-competent tumors.

## Acknowledgements

This work was supported by the NIH Intramural Research Program (NCI ZIA BC 011793). The contributions of the NIH author(s) were made as part of their official duties as NIH federal employees, are in compliance with agency policy requirements, and are considered Works of the United States Government. However, the findings and conclusions presented in this paper are those of the author(s) and do not necessarily reflect the views of the NIH or the U.S. Department of Health and Human Services.

## Conflict of Interest

AT received grants to the institution from Boundlessbio, Gilead, and EMD Serono. AM-9022 was supplied by Volastra under an MTA. The other authors declare no competing interests.

## Supplementary Figures

**Supplementary Figure S1.**
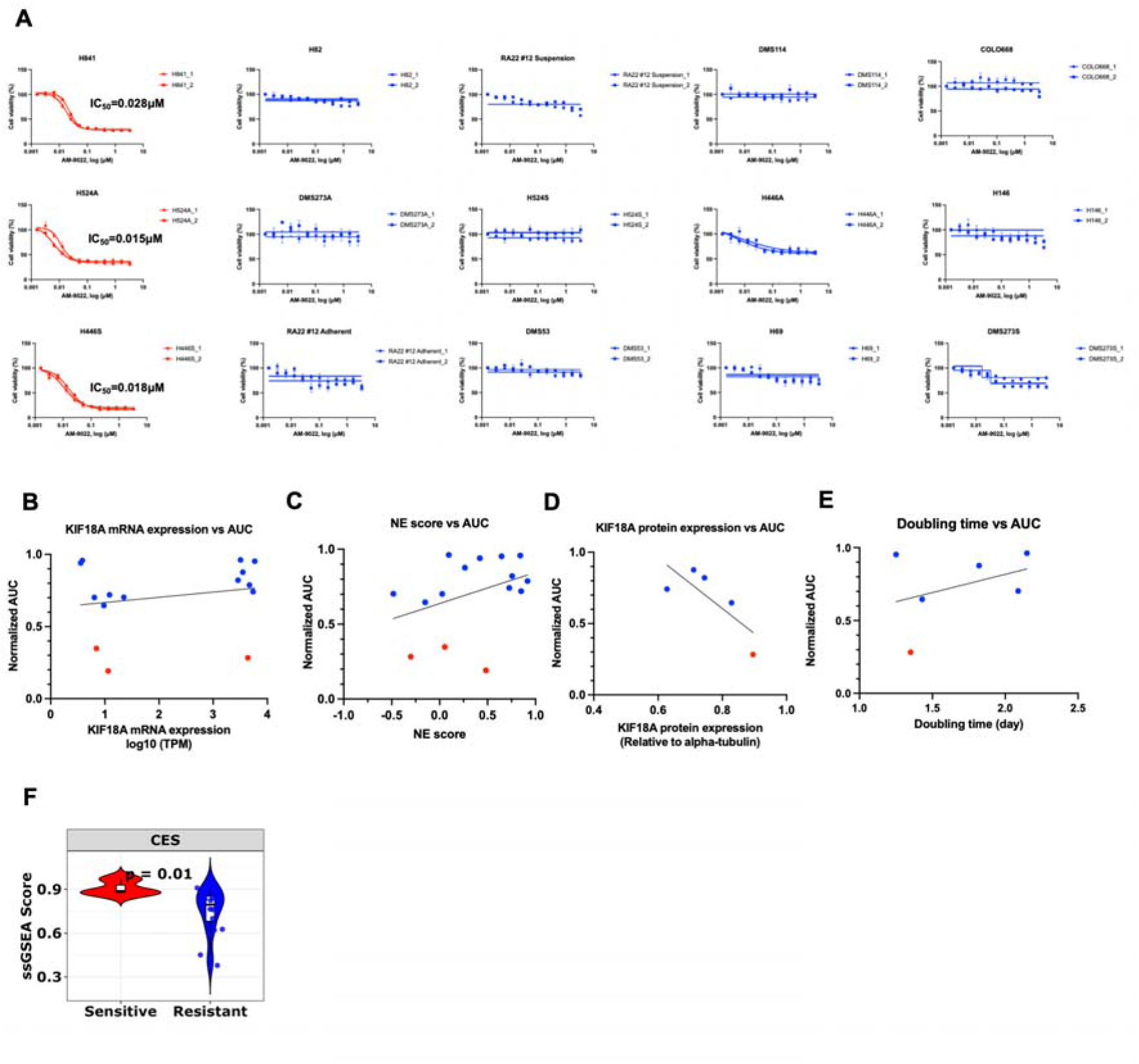
Supplementary analyses supporting the stratification of SCLC cell lines by sensitivity to KIF18A inhibition. **A**, Individual ATP-based dose–response curves for the 15 SCLC cell lines shown in Fig. 2D. For each cell line, two independent experiments are shown, each performed in triplicate, resulting in two dose–response curves per cell line. IC□□ values were calculated for each experiment and were used to classify cell lines as KIF18A inhibitor–sensitive (<0.03 μM) or –resistant (>3 μM), as summarized in Fig. 2D. For clarity, a single representative dose–response curve per cell line is shown in Fig. 2D. **B**, Correlation analysis showing the relationship between normalized area under the curve (AUC) values for AM-9022 sensitivity and KIF18A mRNA expression across a panel of 15 SCLC cell lines. No significant correlation was observed. **C**, Correlation analysis showing the relationship between normalized AUC values and neuroendocrine (NE) scores across 15 SCLC cell lines. No significant correlation was observed. **D**, Correlation analysis showing the relationship between normalized AUC values and KIF18A protein expression across five SCLC cell lines. No significant correlation was observed. **E**, Correlation analysis showing the relationship between cell doubling times and AM-9022 normalized AUC values across six SCLC cell lines. No significant correlation was observed, indicating that KIF18A inhibitor sensitivity is independent of cell proliferation rate. **F**, Violin plots showing single-sample GSEA (ssGSEA) scores for the CES signatures across sensitive and resistant SCLC cell lines. Sensitive exhibited significantly higher CES scores (unpaired t-test, P = 0.01), consistent with elevated chromosomal instability.

**Supplementary Figure S2.**
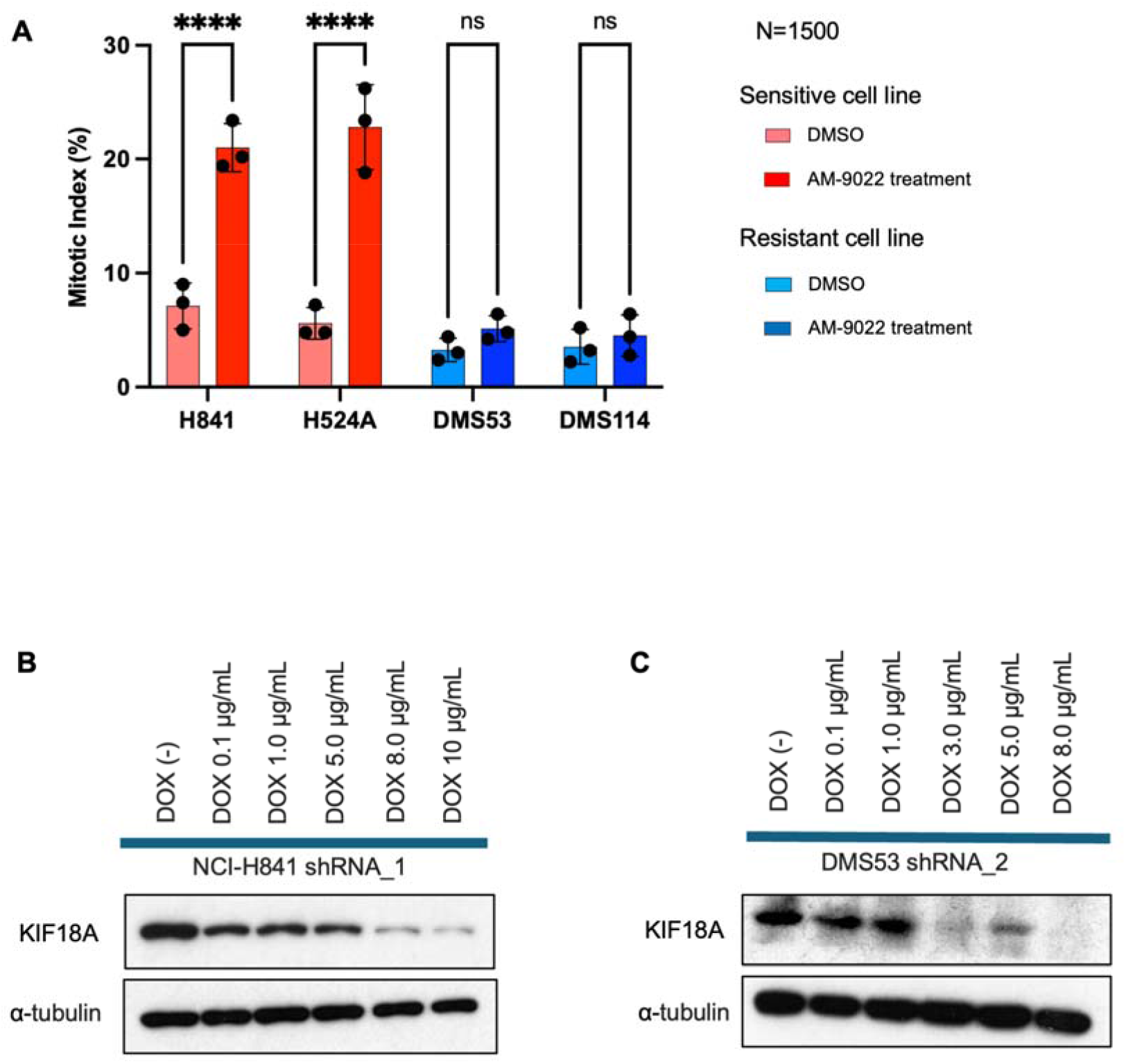
Mitotic index following pharmacologic KIF18A inhibition and validation of doxycycline-inducible KIF18A knockdown in SCLC cells. **A**, Quantification of mitotic index in H841, H524A, DMS53, and DMS114 cells. Cells were treated with DMSO or AM-9022. 1500 cells were counted for each cell line and for each condition across three biological replicates. Dots indicate individual biological replicates (n = 3). Data are presented as mean ± SD. P values were calculated using one-way ANOVA with Tukey’s test. ****, P < 0.0001. **B**, Immunoblot analysis showing KIF18A protein levels in whole cell extracts of NCI-H841 cells treated with different concentrations of DOX to induce KIF18A shRNA. α-Tubulin was used as a loading control. **C**, Immunoblot analysis showing KIF18A protein levels in whole cell extracts of DMS53 treated with different concentrations of DOX to induce KIF18A shRNA_2. α-Tubulin was used as a loading control.

**Supplementary Figure S3.**
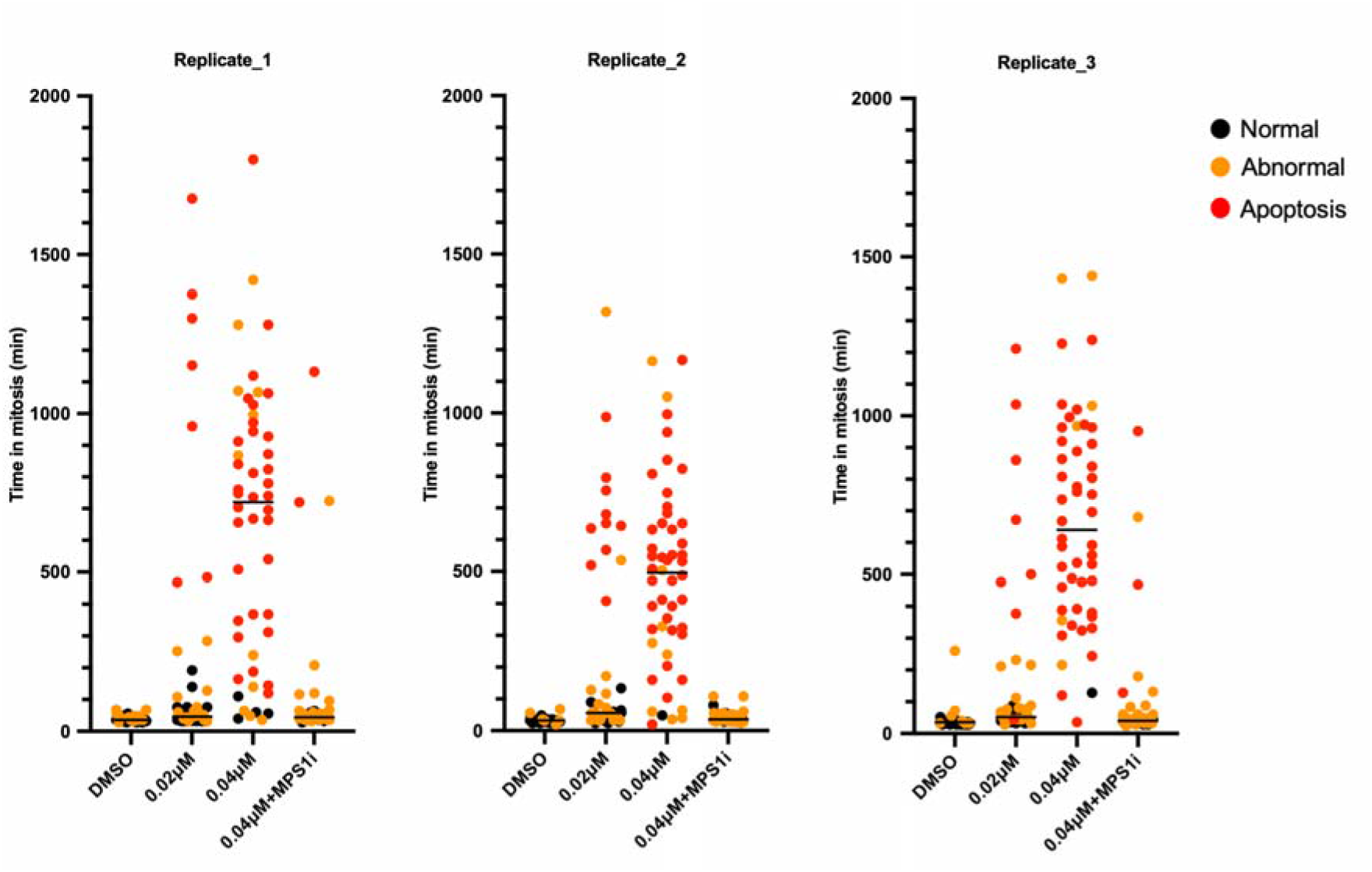
Single-cell analysis of mitotic timing and fate following KIF18A inhibition in NCI-H841 cells. Quantification of mitotic timing in NCI-H841 cells under the indicated treatment conditions. Each dot represents a single cell measured by time-lapse imaging from nuclear envelope breakdown to either anaphase onset or cell death. For each condition, 50 mitotic cells were analyzed and classified based on mitotic outcome as normal mitotic exit, abnormal mitotic exit, or apoptosis.

**Supplementary Figure S4.**
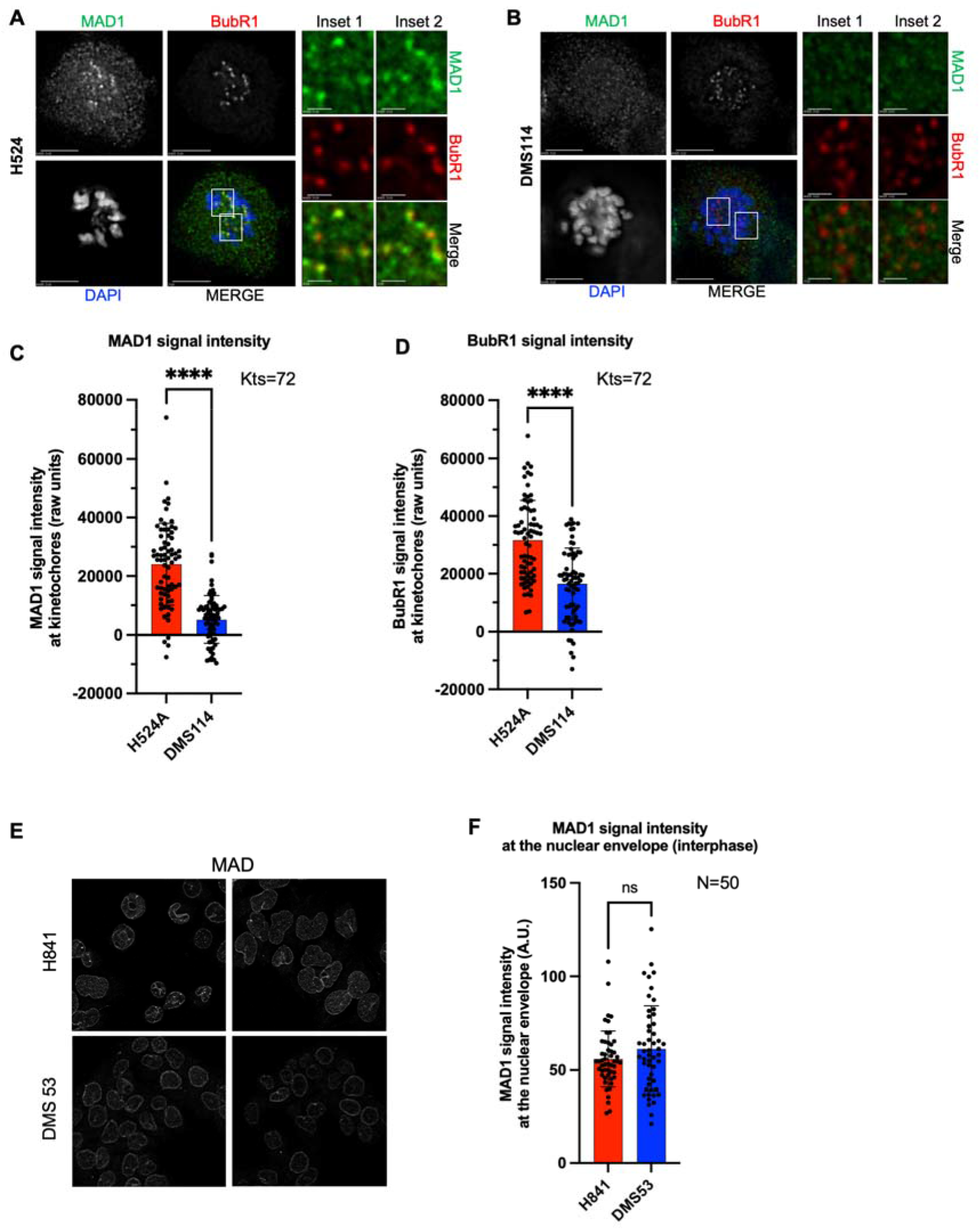
Extended analysis of kinetochore and nuclear envelope localization of spindle assembly checkpoint components in SCLC cells. **A**, Representative immunofluorescence images showing localization of MAD1 (green) and BubR1 (red) at prometaphase kinetochores in monastrol treated NCI-H524A cellss. Insets show zoomed images from white boxed areas in main images. Scale bar: 5 μm for main images and 1 μm for insets. H524A refers to an adherent variant of the NCI-H524 cell line used in this study. **B**, Representative immunofluorescence images showing localization MAD1 (green) and BubR1 (red) at prometaphase kinetochores in monastrol treated DMS114 cells. Insets show zoomed images from white boxed areas in main images. Scale bar: 5 μm for main images and 1 μm for insets. **C**, Quantification of prometaphase kinetochore-associated MAD1 signal intensities in monastrol treated NCI-H524A and DMS114 cells. Bar represents mean ± SD across kinetochores (n=69 for NCI-H524A and n=72 for DM114). Kinetochores from at least 6 cells from two independent experiments were analyzed. Each dot represents a single kinetochore. **D**, Quantification of prometaphase kinetochore-associated BubR1 signal intensities in NCI-H524A and DMS114 cells. Bar represents mean ± SD across kinetochores (n=69 for NCI-H524A and n=72 for DM114). Kinetochores from at least 6 cells from two independent experiments were analyzed. Each dot represents a single kinetochore. **E**, Representative projected immunofluorescence images showing localization of nuclear membrane associated MAD1 in interphase NCI-H841 and DMS53 cells. **F**, Quantification of nuclear membrane-associated MAD1 signal intensities in NCI-H841 and DMS53 cells. Bar represents mean ± SD across average cellular intensity in at least 50 cells from two independent experiments. Each dot represents an average intensity of MAD1 in each cell. P values in panels C, D and F were calculated using Welch’s test. ****, P < 0.0001.

**Supplementary Figure S5.**
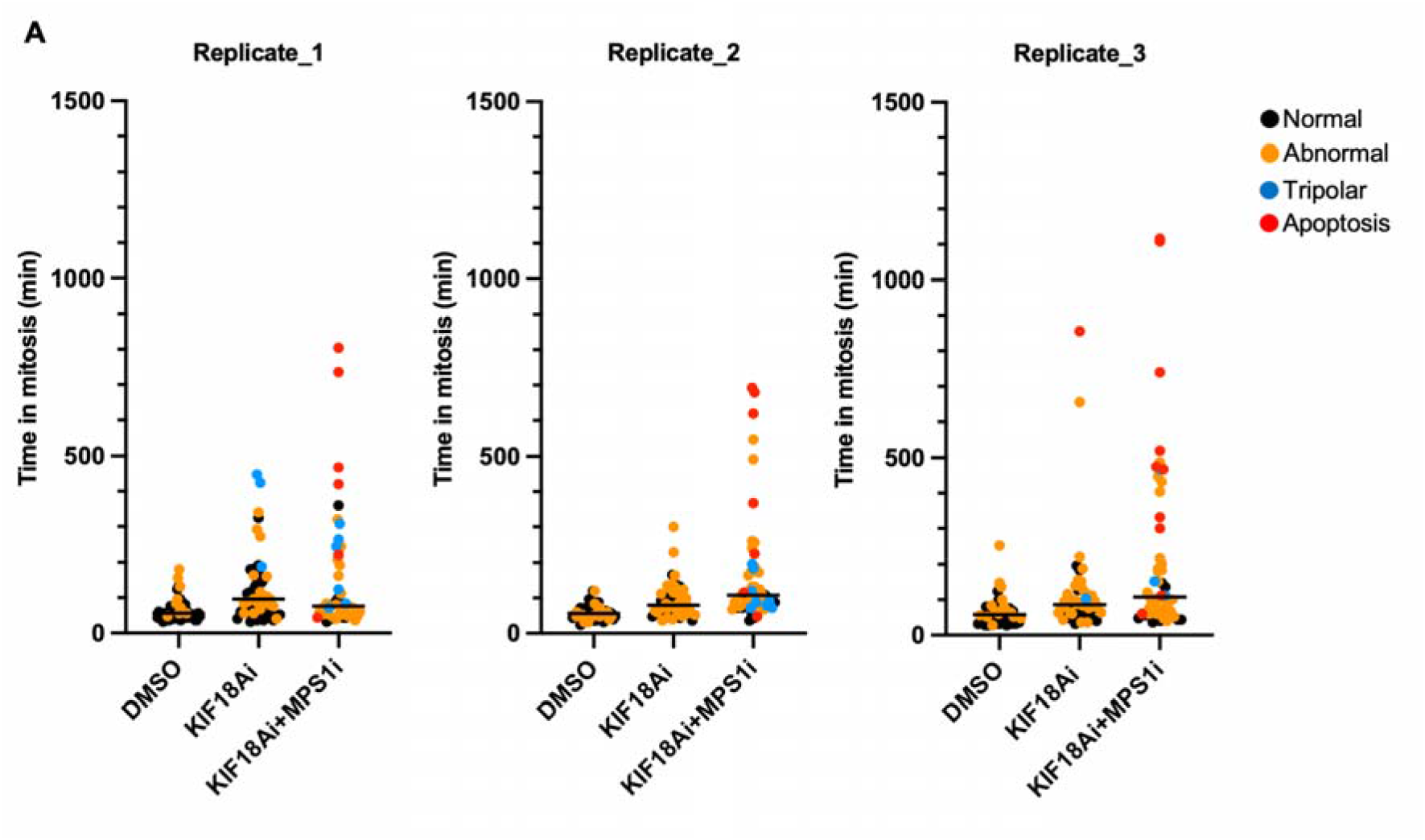
Single-cell analysis of mitotic timing and fate in DMS53 cells following KIF18A inhibition. Quantification of mitotic timing in DMS53 cells under the indicated treatment conditions. Each dot represents a single cell measured by time-lapse imaging from nuclear envelope breakdown to either anaphase onset or cell death. For each condition, 50 mitotic cells were analyzed and classified based on mitotic outcome as normal mitotic exit, abnormal mitotic exit, tripolar, or apoptosis.

## Supplementary Tables

**Supplementary Table 1.**
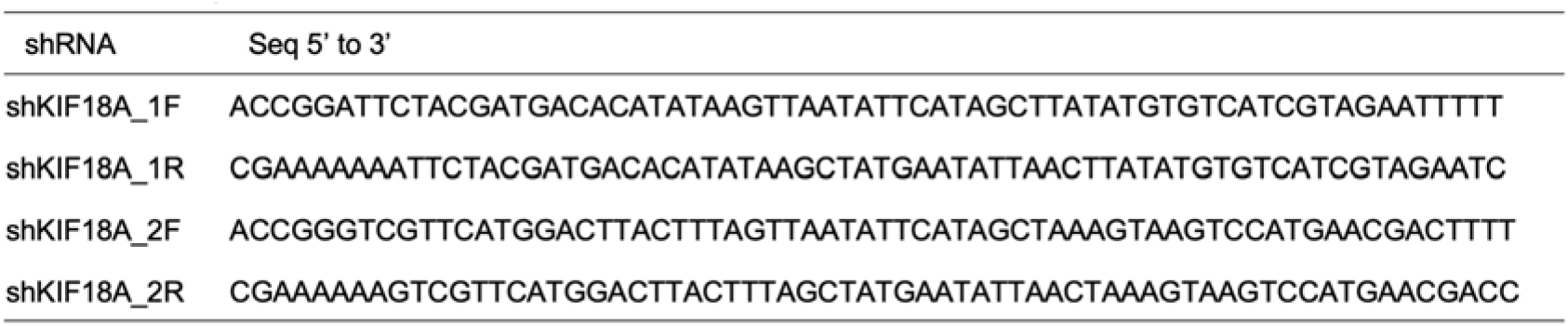
Sequences of shRNAs targeting KIF18A used in this study.

**Supplementary Table 2.** KIF18A expression is shown as log10-transformed transcripts per million (TPM) values derived from RNA-sequencing data. CES scores represent centromere and kinetochore gene expression signature scores calculated from RNA-seq data as gene-level z-scores using a previously defined signature. Replication stress (RepStress) scores and neuroendocrine (NE) scores were obtained from the CellMiner database, as described previously. NE status was assigned based on NE score thresholds. Subtype classification (ASCL1, NEUROD1, POU2F3, or YAP1) was determined based on established molecular criteria.

| Cell line | KIF18A expression<br>log10 (TPM) | CES score | Repstress<br>score | NE score | NE status | Subtype |
| --- | --- | --- | --- | --- | --- | --- |
| NCI-H841 | 3.64 | 0.87 | 1.18 | -0.30 | Non-NE | YAP1 |
| NCI-H446_S | 1.06 | 0.97 | 1.13 | 0.48 | NE | NEUROD1 |
| NCI-H524_A | 0.84 | 0.89 | 1.14 | 0.05 | NE | NEUROD1 |
| DMS 53 | 3.55 | 0.38 | 1.12 | 0.26 | NE | ASCL1 |
| NCI-H69 | 3.74 | 0.70 | 1.15 | 0.73 | NE | ASCL1 |
| NCI-H82 | 3.46 | 0.77 | 1.38 | 0.75 | NE | NEUROD1 |
| DMS 114 | 3.50 | 0.62 | 1.09 | 0.10 | NE | YAP1 |
| NCI-H146 | 3.67 | 0.63 | 1.22 | 0.92 | NE | ASCL1 |
| COLO668 | 3.77 | 0.45 | 1.01 | 0.65 | NE | ASCL1 |
| NCI-H446_A | 0.98 | 0.89 | 1.16 | -0.15 | Non-NE | YAP1 |
| NCI-H524_S | 0.58 | 0.76 | 1.17 | 0.84 | NE | NEUROD1 |
| DMS 273_A | 0.55 | 0.76 | 1.21 | 0.42 | NE | NEUROD1 |
| DMS 273_S | 0.81 | 0.82 | 1.33 | 0.03 | Non-NE | NEUROD1 |
| RA22#12_A | 1.35 | 0.84 | 1.05 | -0.48 | Non-NE | YAP1 |
| RA22#12_S | 1.09 | 0.92 | 1.20 | 0.85 | NE | ASCL1 |

**Supplementary Table 3.** Doubling time is expressed in days. Proliferation rate is expressed as the percentage increase in cell number per day (%/day).

| Cell line | Doubling time<br>(day) | Proliferation rate<br>(%/day) | AM-9022<br>Sensitivity |
| --- | --- | --- | --- |
| DMS114 | 2.15 | 37.86 | Resistant |
| COLO668 | 1.25 | 73.65 | Resistant |
| NCI-H446A | 1.43 | 62.29 | Resistant |
| RA22#12A | 2.09 | 39.23 | Resistant |
| NCI-H841 | 1.35 | 66.61 | Sensitive |
| DMS53 | 1.82 | 46.23 | Resistant |

## Notes

### Competing Interest Statement

Anish Thomas received grants to the institution from Boundlessbio, Gilead, and EMD Serono. AM-9022 was supplied by Volastra under an MTA. The other authors declare no competing interests.

